# Nanofibrous PEDOT-Carbon Composite on Flexible Probes for Soft Neural Interfacing

**DOI:** 10.1101/2021.08.03.454873

**Authors:** Venkata Suresh Vajrala, Valentin Saunier, Lionel G Nowak, Emmanuel Flahaut, Christian Bergaud, Ali Maziz

**Affiliations:** Laboratory for Analysis and Architecture of Systems (LAAS), CNRS, Toulouse, France; Centre de Recherche Cerveau et Cognition (CerCo), CNRS, Toulouse, France; CIRIMAT, Université de Toulouse, CNRS, route de Narbonne, F-31062 Toulouse, France

**Author notes:** Corresponding authors: Dr. Ali MAZIZ, LAAS-CNRS.

**Keywords:** PEDOT-Carbon, Carbon nanofibers, Porous electrode composite, Flexible neural interfaces, Electrophysiological recording, Neural stimulation

## Abstract

In this study, we report a flexible implantable 4-channel microelectrode probe coated with highly porous and robust nanocomposite of poly(3,4-ethylenedioxythiophene) (PEDOT) and carbon nanofiber (CNF) as a solid doping template for high-performance in vivo neuronal recording and stimulation. A simple yet well-controlled deposition strategy was developed via in situ electrochemical polymerization technique to create a porous network of PEDOT and CNFs on a flexible 4-channel gold microelectrode probe. Different morphological and electrochemical characterizations showed that they exhibit remarkable and superior electrochemical properties, yielding microelectrodes combining high surface area, low impedance (16.8 ± 2 MΩ.μm^2^ at 1 kHz) and elevated charge injection capabilities (7.6 ± 1.3 mC/cm^2^) that exceed those of pure and composite PEDOT layers. In addition, the PEDOT-CNF composite electrode exhibited extended biphasic charge cycle endurance, resulting in a negligible physical delamination or degradation for long periods of electrical stimulation. In vitro testing on mouse brain slices showed that they can record spontaneous oscillatory field potentials as well as single-unit action potentials and allow to safely deliver electrical stimulation for evoking field potentials. The combined superior electrical properties, durability and 3D microstructure topology of the PEDOT-CNF composite electrodes demonstrate outstanding potential for developing future neural surface interfacing applications.

## 1. Introduction

Neural electrodes provide the critical interface between the nervous system and electronics. Well-defined anatomical regions from the brain can be the targets of implanted microelectrodes, enabling localized neuromodulation by either recording or delivering electrical signals at the level of individual neuron [1–3]. Such capabilities have been critically important for supporting neuroscience research along with emerging clinical devices aimed at treating debilitating disorders, including deafness [4, 5], paralysis [6], blindness [7], Parkinson’s disease [8], epilepsy [9] and other disorders [10, 11]. In all of these applications, the crucial material-dependent problem is developing microelectrode array that sense and/or stimulate neural activity from small, targeted groups of neurons with high fidelity and long-term reliability [12].

Conventional implantable microelectrode arrays, made of silicon backbone and noble metal electrodes, such as gold (Au) platinum (Pt) or iridium (Ir) become routine in animal research and have occasionally been used in humans [13, 14]. However, they are not suitable for long-term use due to limitations regarding the electrical and mechanical mismatches with the surrounding tissue [12, 15]. Decreasing the size of an electrode active site, to ideally target single neuron, results in low capacitance and high impedance at the electrode/tissue interface, which seriously impacts recording resolution and stimulation capabilities [16–18]. For neuronal recording, the electrode impedance contributes to the noise, and high impedance electrodes are expected to have a low signal-to-noise ratio (SNR) [19, 20]. For neuronal stimulation, an ideal electrode should display a high storage capability to safely inject current pulses with minimal potential transients at the electrode/tissue interface, thus decreasing both electrode polarization and heat accumulation during stimulation [18, 20, 21].

Besides, immune reaction occurs through the mechanical mismatch between rigid electrodes and the neural tissue, triggering inflammatory responses and glial scar formation, which may lead to encapsulation of the electrodes and subsequent device failure [15, 22]. These device failures appear in the form of electrical recording degradations including increased impedance, increased noise levels and decreased signal amplitudes [22]. In this regard, there has been a demand to fabricate electrodes on flexible substrates that, by showing smaller hardness mismatch, provide a more adaptable interface to neural tissue. In addition, the electrode material deposited on flexible substrate should display low electrical impedance and high charge-transfer capacity without substantially increasing the site geometric surface area [23].

To tackle this challenge, various types of organic electroactive materials have been employed such as conductive polymers (CPs) [24–26], carbon-based nanomaterials *i.e.* carbon nanotubes (CNTs) [27, 28], graphene [29], reduced graphene oxide (r-GO) [30] and their nanocomposites [31–33], to create much desired porosity and softness at the electrode/tissue interface. Among them, the conducting polymer Poly (3,4-ethylenedioxythiophene) (PEDOT) has been a popular choice due to its mixed electronic and ionic conductivities, high-quality electrochemical performances, together with excellent biocompatibility, softness, and ease of functionalization [24, 34–37]. Several reports showed that PEDOT coatings, doped with different counter ions such as poly(styrene sulfate) (PSS) [38, 39], Nafion [40], tosylate [41, 42], dodecyl sulfate [43] or ClO_4_^-^[44], can significantly decrease the electrode impedance (30-250 MΩ/ μm^2^) and increase the charge-injection capacity (1-3 mC/ cm^2^) as compared to flat metal sites of similar geometric area [18, 45–47]. In addition, *In vitro* studies have demonstrated that PEDOT coating would also present a good substrate for the growth of various cell types in biological and tissue engineering areas, wherein PEDOT directly interacts with cells or tissues [48, 49]. Despite its promising outlook, PEDOT is yet to be perfected as a coating material for neural electrodes, especially in terms of extended charge injection capabilities and long-term adhesion stability [50, 51]. For example, brain stimulation applications demand high and stable charge injection capabilities, where the electrode materials should sustain few thousands to millions of cycles of electrical stimulations pulses without corrosion, tissue damage, or delamination [20].

Multiple strategies have been suggested to reinforce PEDOT coating, either by modifying the monomer itself [52], using adhesion promoters [53], or by the incorporation of charged carbonaceous nanomaterials [31, 54]. In previous studies, composite materials made up of CNTs[31, 55] or r-GO [30], in combination with PEDOT, have been coated on metal microelectrodes to decrease the impedance and ramp-up the charge injection limits [56], even beyond the PEDOT: PSS capabilities for both acute and chronic stimulation tests [57, 58]. It was also reported that these mechanically strong carbonaceous materials function as reinforcing elements within the composite, preventing the PEDOT film from undergoing deformation and cracking during prolonged redox reactions [54]. This excellent performance makes the combination of PEDOT with carbon-based nanomaterials a highly promising candidate material for the development of long-lasting neural interfaces. However, as of now, most of the research has focused on either the selection of near perfect electrode material with superior electrical properties or controlled 3D surface macro porous pattering of electrode to promote an intimate contact with the neural tissue. In contrast, less attention has been paid to the combination of both.

Recently, we have demonstrated the feasibility of a novel composite material by combining PEDOT with carbon nanofibers (CNFs) through a simple and reproducible electrodeposition method [32]. CNFs exhibit extraordinary strength, high modulus of elasticity (940 GPa), and provide an extremely large surface area for charge transfer and cell attachment [59]. Since, they contain basal graphite planes and edge planes, upon oxidation, their outer surface can be electrochemically functionalized with PEDOT molecules. In addition, as a result of their extremely high edge proportion with very high aspect ratios and inherent herringbone morphology, making them excellent nanoscale building block to establish interconnected, three-dimensional (3D) macroporous structures, in combination with PEDOT. We have recently shown that the combination of CNFs and PEDOT on rigid microelectrode array (MEA) resulted in as strong synergetic effect between the two components in the single composite leading to remarkable electrochemical properties, as well as a reliable *in vitro* neurotransmitter monitoring using amperometric techniques [32]. These results suggest PEDOT-CNF composites as a most interesting electrode material for applications in neuroprostheses and neurophysiology research. In this context, to go further into the development of neural interfacing devices for *in vitro* and *in vivo* applications, we are reporting a method for preparing macroporous, stable and electrically superior PEDOT, using CNFs as a solid dopant template, on ultra-flexible penetrating neural microelectrodes. We developed a well-controlled one-shot deposition method and optimized it for preparing macroporous PEDOT-CNFs nanocomposite via *in situ* electrochemical polymerization technique on flexible parylene-based neural probes. This flexible substrate- provides the means to decrease the mechanical mismatch at the electrode/tissue interface. We showed that PEDOT-CNF hybrid neural microelectrodes exhibit remarkable electrochemical properties, yielding microelectrodes combining low impedance, high surface area, and elevated charge injection capabilities. This device was further tested for neural recording and stimulation in the hippocampus of mouse brain slice in vitro. The obtained results opened great prospects for the development of next-generation microelectrodes for applications in brain therapies.

## 2. Materials and Methods

### 2.1. Fabrication of the neural implant and device packaging

The photomask designs of the flexible parylene C-based probe, with four micro disk electrodes, are inspired from our previous works [38, 60]. The fabrication procedure is schematically illustrated in Fig. 1a. A 23 μm thick film of Parylene C was deposited using chemical vapour deposition (Comelec C-30-S at 700 °C) on 4-inch SiP wafer. Next 50 nm thick Ti and 200 nm thick Au layers were deposited and patterned, using a conventional physical vapour deposition technique followed by a lift-off with AZ-nLof 2035 (Micro chemicals). Subsequently, 1.3 μm thick parylene C was deposited as a passivation layer, followed by an annealing step at 110°C for 16 hours under nitrogen flow to increase the adhesion between parylene C-gold sandwich layers. Next, the electrode surfaces (40 μm diameter) and the corresponding connection pads were realized by, photo-patterning with 5 μm thick AZ4562 photoresist, followed by the etching of the thin parylene C passivation layer using O_2_ plasma reactive ion etching ICP-RIE (Trikon Omega 201). Later, the implants were anistropically etched to establish smooth outlines and vertical sidewalls, using 50 μm thick BPN photoresist (Intervia BPN-65A, Dupont) and a deep reactive ion etching (ICP-DRIE) step. After that, the implants were peeled-off carefully by placing the entire wafer into DI water for at least 2 hours. Later, the released implants were stripped off from the leftover photoresist using TechniStrip NF52 (Microchemicals), thoroughly washed in DI water and stored in a dry place. Finally, the implants were bonded to a customized flexible ribbon cable with golden traces (AXO-00021, pro-POWER, China) by using epoxy silver and photosensitive glue.

**Fig. 1:**
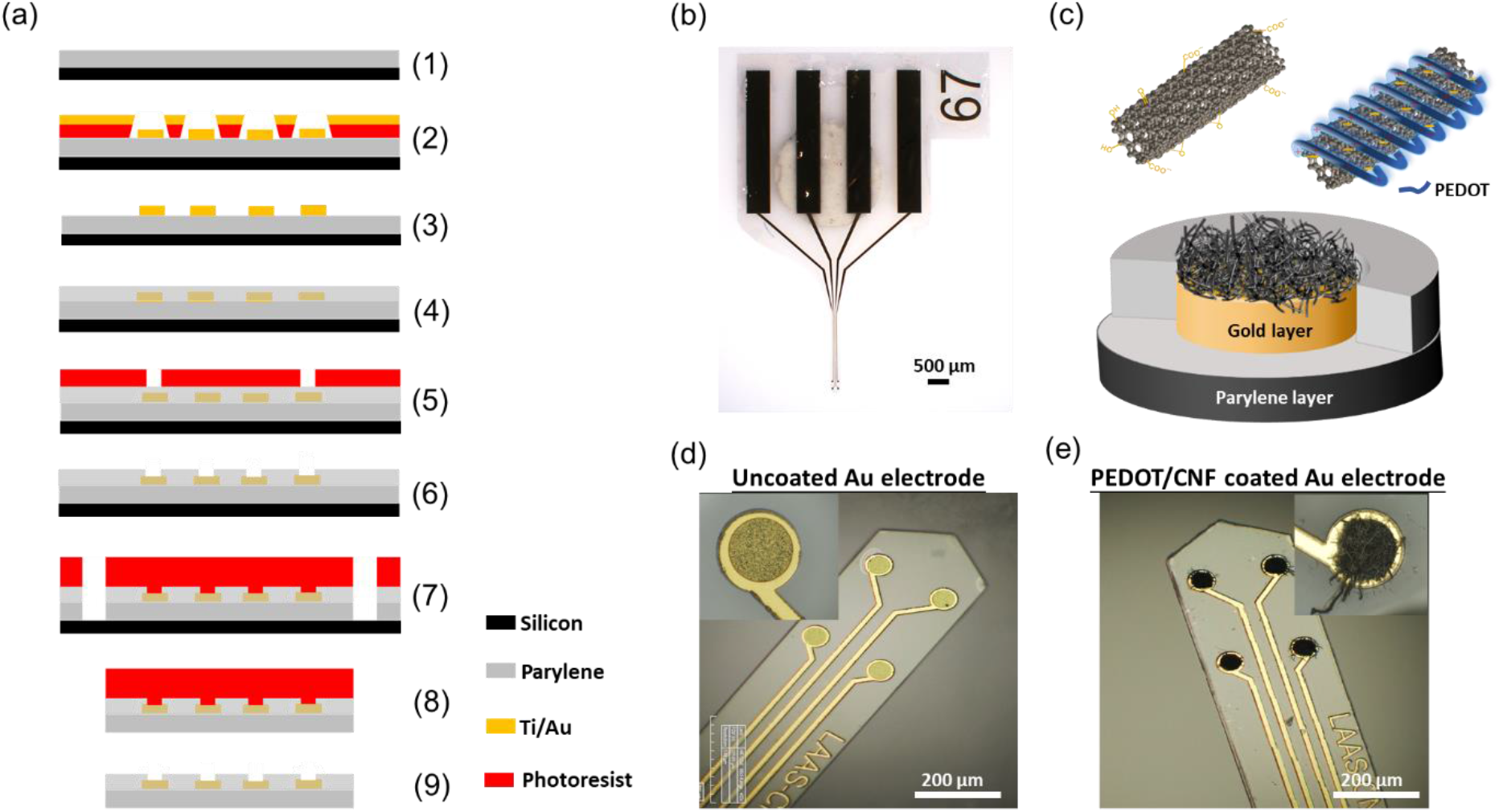
Schematic illustration and optical images of neural implant. (a) Schematic overview of steps involved in the microfabrication procedure- (1) 23 μm thick Parylene C deposition on SiP wafer; (2) Photo-patterning of nLof and gold layer deposition (Ti/Au-50/200 nm); (3) nLof removal; (4) 2^nd^ Parylene C layer (1.3 μm) deposition and anneal at 110°C for 16h; (5) Spin coating of 5μm thick AZ4562 photoresist; (6) RIE with ICP-RIE oxygen plasma; (7) Photo-patterning with BPN photoresist and RIE with ICP-DRIE oxygen plasma; (8) Stripping of implants from SiP wafer and (9) Development with NF52 and cleaning with DI water. (b) Optical micrograph of the neural implant. (c) Schematic representation of the PEDOT-CNF composite deposition. (d) and (e) Optical micrographs showing the neural microelectrodes before and after surface modification with PEDOT-CNF composite.

### 2.2. Functionalization of Carbon nanofibers (CNFs)

Raw CNFs (Pyrograf^®^-III, PR-19-XT-PS, pyrolytically stripped, platelets conical, >98% carbon basis, 20-200 μm) were oxidized using wet chemical oxidation process where 300 mg of CNFs were placed in 200 mL of the oxidizing solution (15 M) HNO_3_/ (18 M) H_2_SO_4_ and sonicated for 15 min, followed by 2 hours of reflux at 70 °C [61, 62]. This process helps to remove impurities (metallic particles and amorphous carbon) from the sample and make them hydrophilic, so that the CNFs can be dispersed in water [63]. Following the acid treatment, the CNF dispersion was washed with DI water to reach a neutral pH, and stored at 4°C.

### 2.3. Electrochemical deposition of PEDOT-CNF composite

The stock solution of oxidized CNFs was first vortexed for 10 minutes and homogenized by sonication. Next, the sample was dispersed in DI water, at a concentration of 1 mg/ml, along with 10 mM EDOT (Sigma Aldrich). Later, the CNFs-EDOT mixed suspension was incubated under vortex for 2 days at room temperature. Prior to use, the suspension containing CNFs and EDOT polymer was sonicated for 2 min and vortexed again for 15 min to obtain a homogenized dispersion. The gold microelectrodes were electrochemically cleaned by cycling at 200 mV/s between −0.3 V and 1.4 V vs Ag/AgCl in 0.5 M H_2_SO_4_ using cyclic voltammetry (Fig. S1a). PEDOT-CNF composites were galvanostatically deposited on the clean Au electrode surface by applying the current density of 10 pA/μm^2^ with a range of charge densities (1, 2, 4, 6 and 8 nC/μm^2^) (Fig. 2 and Fig. S1b).

**Fig. 2:**
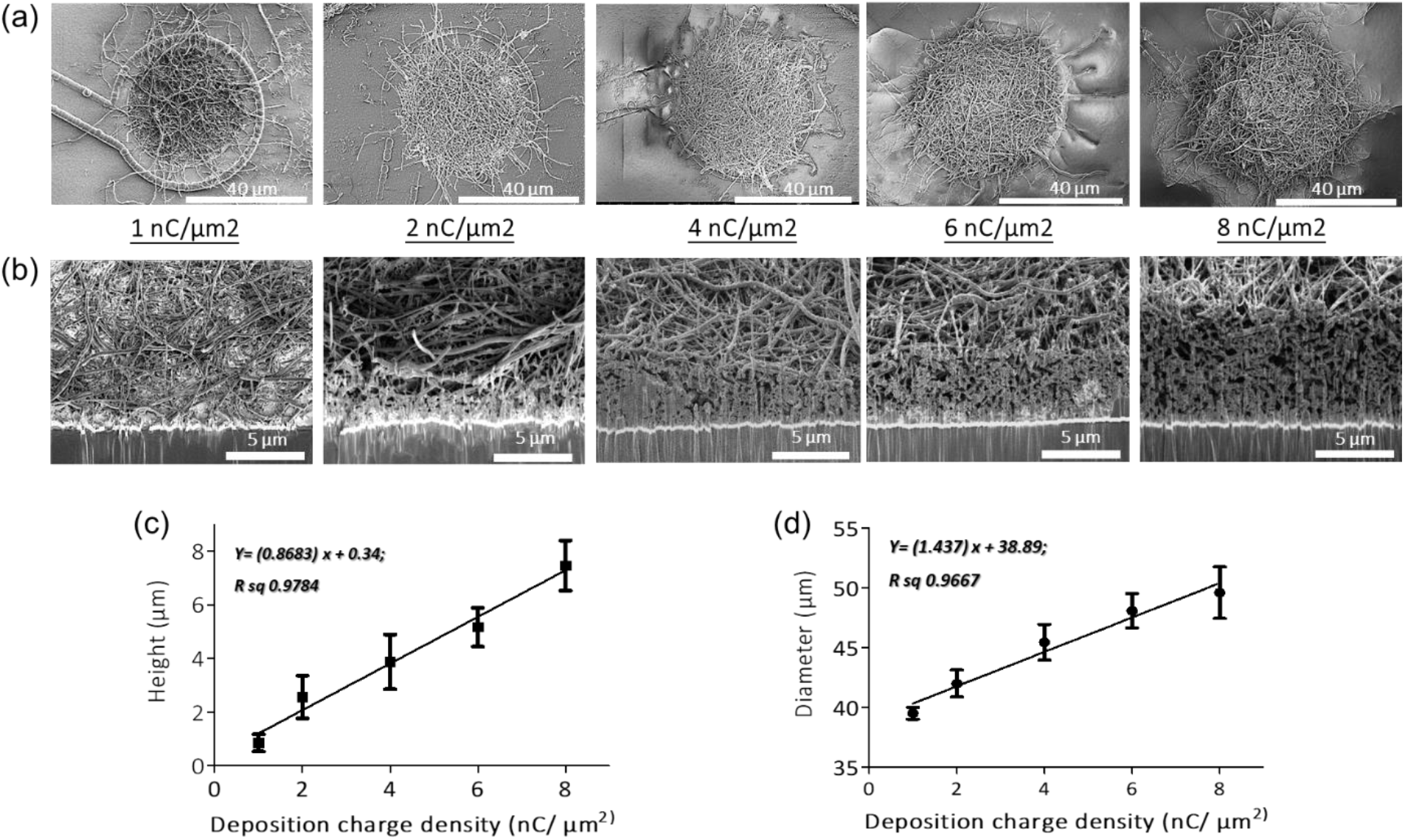
SEM micrographs of (a) top views and (b) cross-sectional views of PEDOT-CNF composites on flexible gold electrodes with different surface charge densities ranging from 1 to 8 nC/μm^2^. PEDOT-CNF films were galvanostatically deposited on gold microelectrodes with a constant current density of 10 pA/μm^2^, using two-electrode configuration. Plots describing the evolution of deposition height (c) and diameter (d) of the composite versus deposition charge density. The error bars represent the standard deviation where n=6.

### 2.4. Optical characterization

Optical microscope images of the implant, before and after the composite deposition (Fig. 1d-e) were obtained with a HIROX microscope (HI-SCOPE Advanced KH-3000). The morphological information of the PEDOT-CNF composite coatings on flexible microelectrode array (Fig. 2) and the corresponding EDX characterization (Fig. 3) were obtained by using FEG Schottky High resolution Helios 600i Dual FIB scanning electron microscope.

**Fig. 3:**
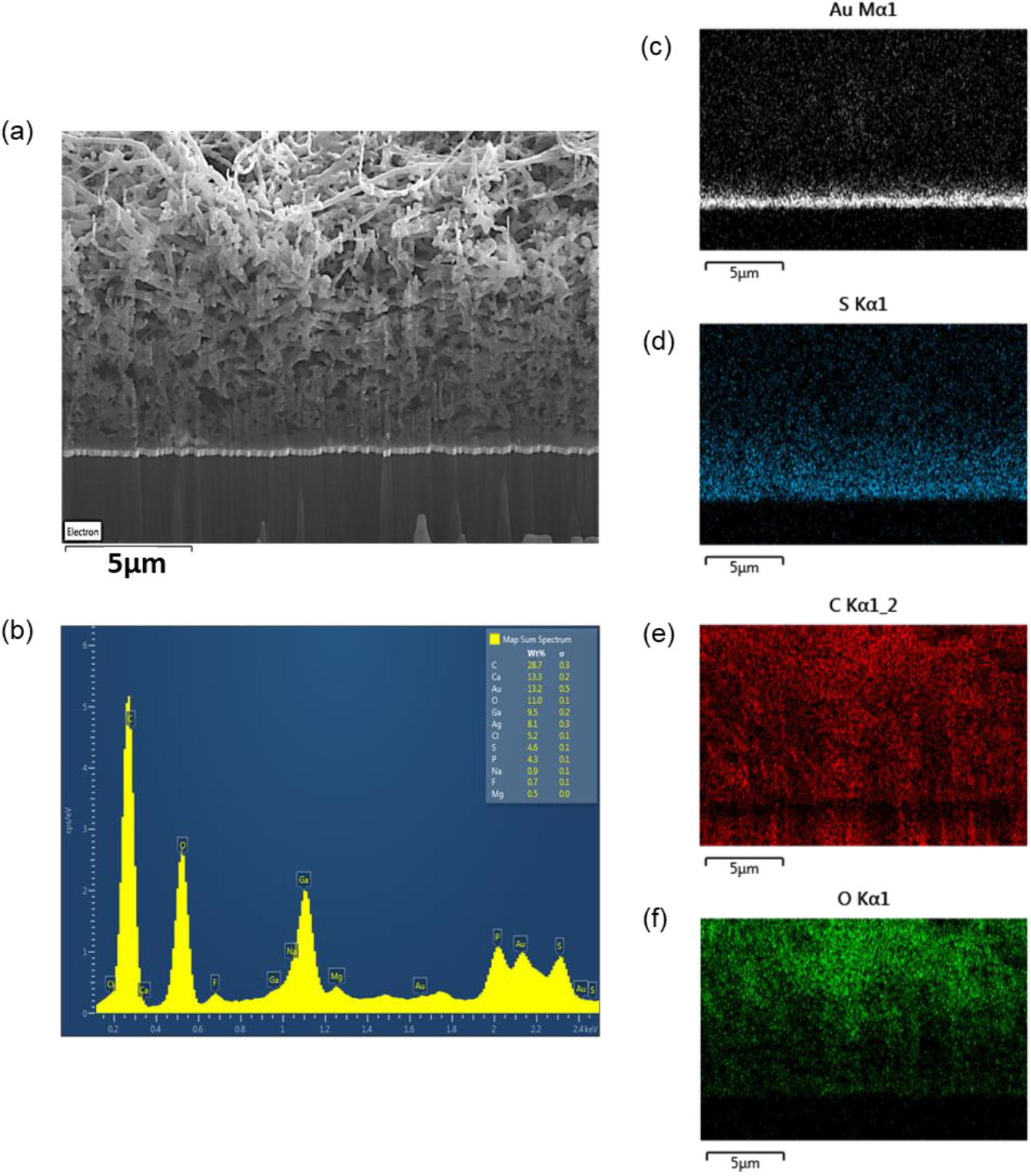
(a) SEM image and (b) EDX spectrum of porous PEDOT-CNF composite (8 nC/μm^2^) deposited on flexible gold electrodes. Elemental mapping images confirm the presence of (c) gold, (d) sulfur, (e) carbon and (f) oxygen in the PEDOT-CNF composite.

### 2.5. Electrochemical characterization

Prior to the characterization, the probes were rinsed with DI water and immersed in artificial cerebrospinal fluid (aCSF) for at least 30 min. For physiological relevance, the aCSF composition mimicked the mammalian ionic CSF composition and consisted of (in mM): NaCl 124, NaHCO_3_ 26, KCl 3.2, MgSO_4_ 1, NaH_2_PO_4_ 0.5, CaCl_2_ 1.1, glucose 10, and bubbled with 95% O_2_ and 5% CO_2_ (pH 7.4) [64]. Probes were proceeded further with charge storage capacity (CSC) (Fig. 4a-b), electrochemical impedance spectroscopy (EIS) (Fig. 4c-f) and charge injection limit (CIL) measurements (Fig. 5 and 6). Electrochemical characterizations were performed in a three-electrode configuration, using a thick (2-mm diameter, ^~^5 mm^2^) Pt wire (WPI, 99.99%) as counter electrode (CE) and an Ag/AgCl reference coil electrode.

**Fig. 4:**
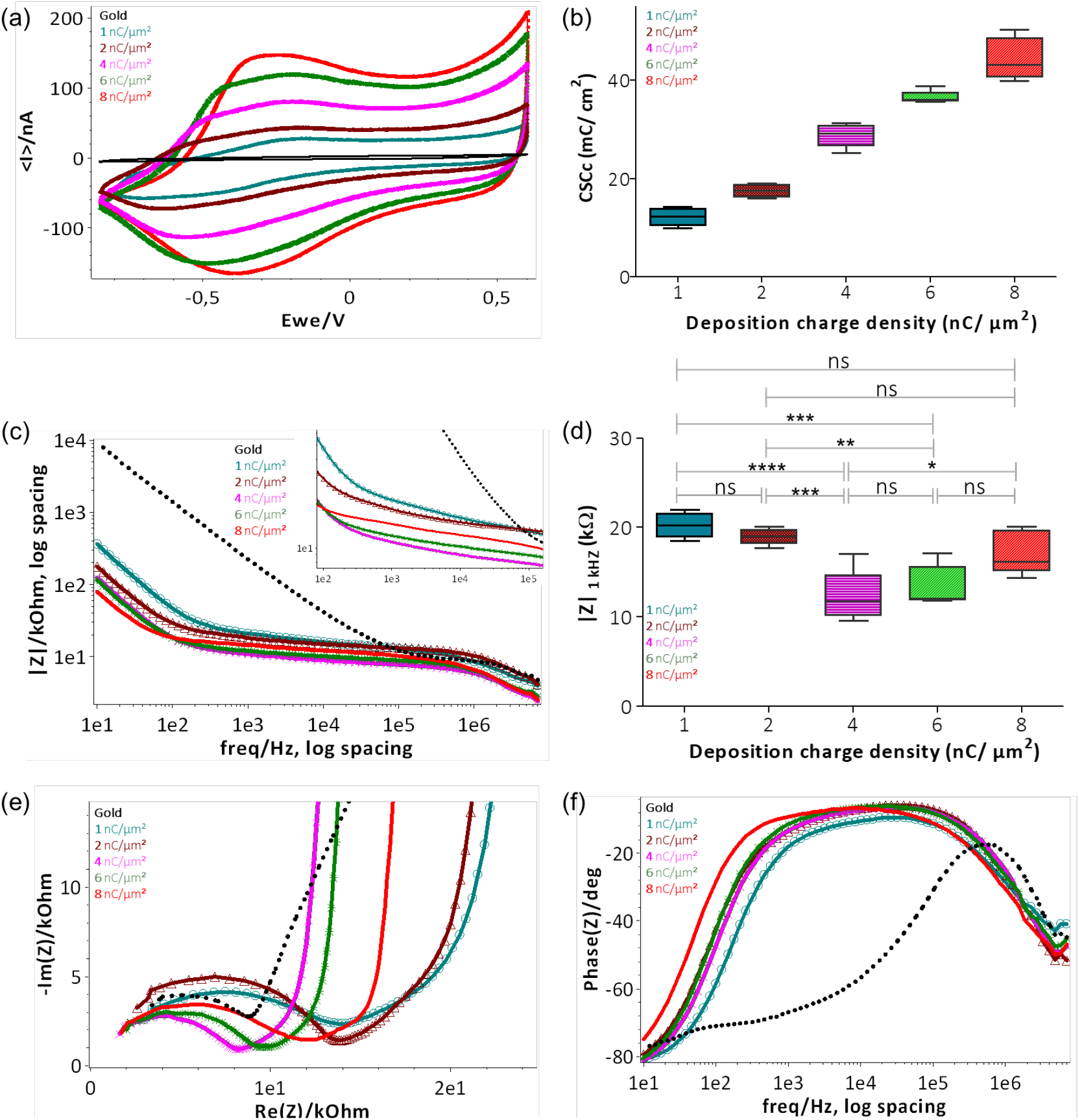
Electrochemical characterization of PEDOT-CNF composite deposited on flexible gold surface with different surface charge densities: (a) CSCc measurements by CV in aCSF at 200 mV/s *vs* Ag/AgCl ref electrode; (b) Plot representing the evolution of CSCc with respect to the deposition charge density; (c) Bode plot representing the |Z| *vs* frequency over a frequency range of 10 Hz to 7 kHz in aCSF at 0V *vs* Ag/AgCl ref electrode; (d) Plot of impedance |Z| _1 kHz_ responses vs deposition charge density. The statistical differences between deposition conditions were assessed by ANOVA followed by Tukey’s posthoc test, where ****, ***, ** and ns represent P<0.0001, P<0.001, P<0.01 and no significant difference, respectively (n=5); (e) Nyquist plots and (f) phase angle measurements obtained by EIS. Deposition charge density conditions were represented in different colors, along with the plain gold electrode (black color) in the above all graphs. (For interpretation of the references to colors in this figure legend, the reader is referred to the web version of this article).

**Fig. 5:**
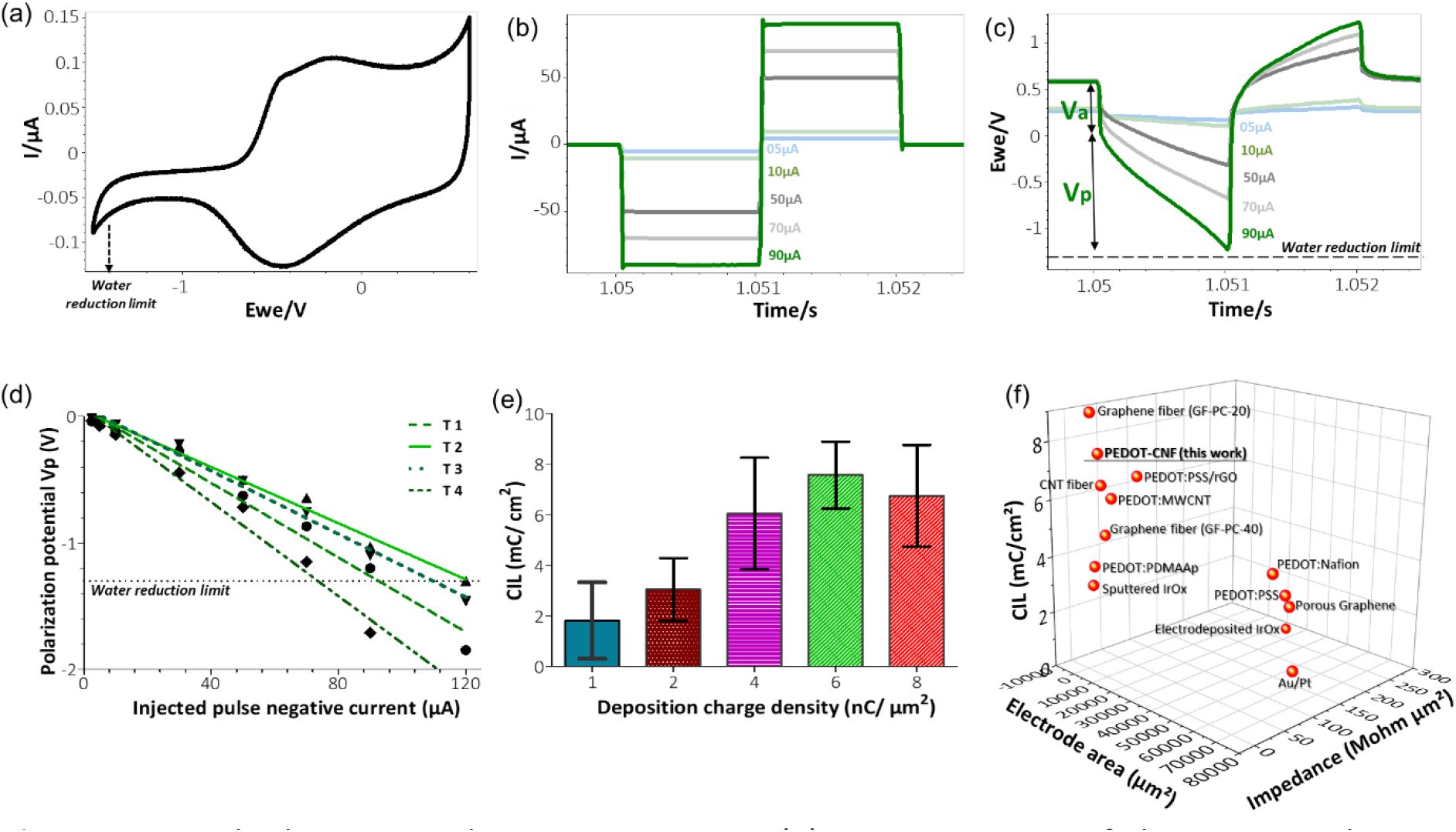
*In vitro* biphasic stimulation assessment: (a) Determination of the water reduction potential by CV of PEDOT-CNF composite in aCSF at 200 mV/s Vs Ag/AgCl. Biphasic charge-balanced current pulses (b) and voltage responses (c) at different charge injections ranging from 5 μA to 90 μA; (d) Polarization potentials (Vp) measured under different current pulse amplitudes. The deposition condition of 6 nC/μm^2^ was considered here and the data corresponding to 4 trials are represented; (e) Evolution of the charge injection limit (CIL) values with respect to the deposition charge density. (f) Comparison of the Impedance, CIL performances and geometric surface area of PEDOT-CNF composite deposited on the flexible gold electrode surface with other flexible neural electrode arrays. The corresponding values and references are reported in Table S1. (For interpretation of the references to colors in this figure legend, the reader is referred to the web version of this article).

**Fig. 6:**
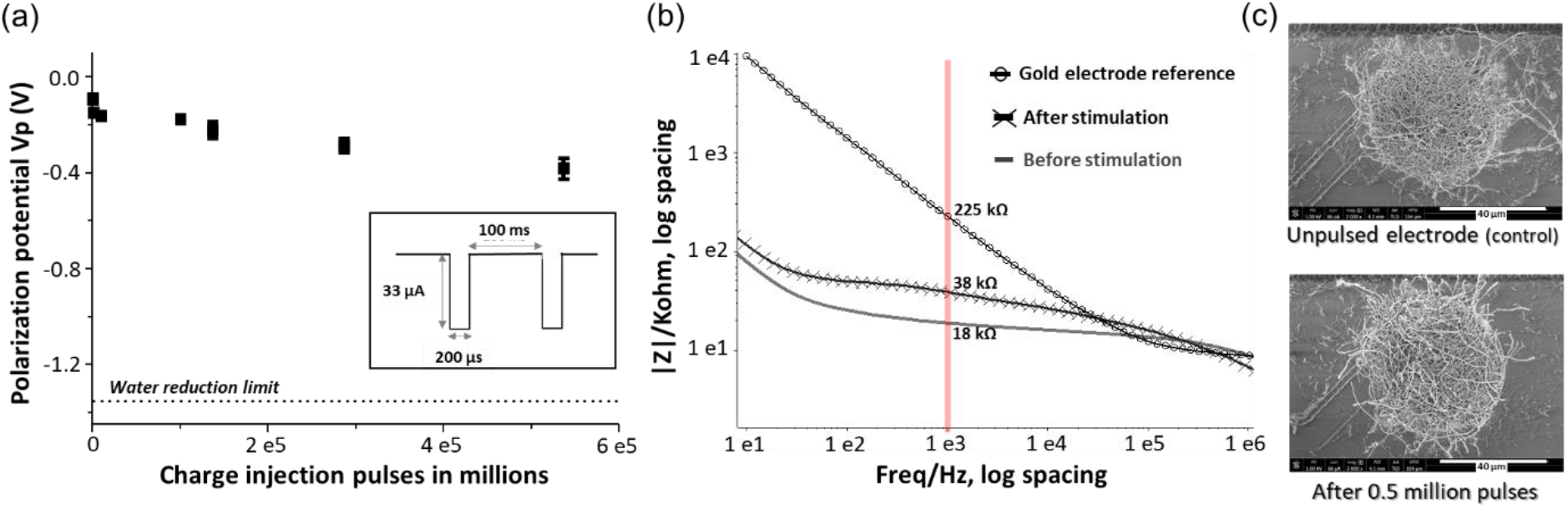
Long term stability assessment of PEDOT-CNF composite: (a) Evolution of polarization potential over 0.5 million charge injection pulses at 0.5 mC/cm^2^ in aCSF. Inset-Schematic representation of the biphasic charge-balanced current waveform having an amplitude of 33 μA and 200 μs pulse duration at 10 Hz. (b) EIS monitoring of PEDOT-CNF composite before and after 0.5 million stimulation pulses. (c) SEM observation of the composite after the stimulation and comparison with the unpulsed electrode (control).

Impedance measurements (EIS) were performed between 10 Hz and 7 MHz, using a 10 mV AC signal at 0 V *vs* Ag/AgCl, whereas the cathodic charge storage capacity measurements (CSCc) were carried out by launching cyclic voltammetry on a low current potentiostat channel (BioLogic VMP3), between 0.6 V and −0.8 V in aCSF at room temperature. Each electrode sample was swept for two cycles and the CSCc was calculated as the time integral of the cathodic current recorded over a potential range of 0.6 V to −0.85 V in the second cycle.

To estimate the charge injection limit, voltage transient measurements were carried out at different input currents by applying charge-balanced biphasic current pulse waveforms at 10 Hz, with pulse durations of 1 ms (Fig. 5) and 200 μs (Fig. 6), using a Bio-Logic VSP3 potentiostat. The negative polarization potential (V_p_) was calculated by subtracting the initial access voltage (V_a_) due to solution resistance from the total voltage (V_max_). The charge injection limits were calculated by multiplying the current amplitude and pulse duration at which the polarization potential reaches the water reduction limit (−1.3 V), divided by the geometric surface area of the electrode.[20]

Long-term stimulation stability testing was assessed by launching at least 0.5 million continuous current pulses of 33 μA at 10 Hz, thereby employing the charge injection capacity of 0.5 mC/cm^2^. The electrochemical differences before and after pulsing were measured with EIS. The electrochemical cell was sealed properly to avoid evaporation of the electrolyte during the measurements.

### 2.6. Brain slice preparation and electrophysiological measurements

All procedures were conducted in accordance with the guidelines from the French Ministry of Agriculture (décret 87/848) and from the European Community (directive 86/609) and was approved by the Ministère de l’Enseignement Supérieur, de la Recherche et de l’Innovation (N° 15226-2018052417151228). Adult (>2-month-old) wild type female mice were anesthetized with isoflurane and killed by decapitation. All following procedures were made in the presence of oxygenated (95% O_2_ and 5% CO_2_) and ice-cold modified, artificial cerebrospinal fluid (maCSF) whose composition was (in mM): NaCl 124, NaHCO_3_ 26, KCl 3.2, MgSO_4_ 1, MgCl_2_ 9, NaH_2_PO_4_ 0.5, and glucose 10 [64]. The upper part of the skull was drilled off and the whole brain was carefully removed and glued on a pedestal for slicing. 400 μm-thick coronal brain slices were cut on a vibratome (752 M vibroslice, Campden Instrument, UK), whose chamber was filled with ice-cold oxygenated maCSF. The slices were kept at room temperature for at least one hour in an *in vivo*-like artificial cerebrospinal fluid (aCSF, composition in “electrochemical characterization” above), aerated with 95% O_2_ and 5% CO_2_ (pH 7.4). For recording and stimulation, a brain slice was fixed on the mesh of a submersion type recording chamber (Scientific System Design, Mississauga, Ontario, Canada), as shown in the Fig. 7a. The recording chamber was continuously supplied in oxygenated aCSF that was gravity fed at a flow rate of 3-3.5 ml/min. The temperature was maintained at 33-34°C. The neural microelectrodes were positioned in the hippocampal regions (CA1 and CA3) of the brain slice, using a 3D micromanipulator. Tungsten-in-epoxy lite microelectrodes (FHC, 0.2–0.3 MΩ) were also used for parallel recording and stimulation. Signals were amplified (final gain: × 10^4^) and filtered with a NeuroLog recording system (Digitimer Ltd, UK) and digitized with a 1401plus interface (CED systems, Cambridge, UK) with a digitization rate of 20 kHz. The signals were visualized online and analysed offline using spike2 software (CED) and custom scripts within Spike2 software.

**Fig. 7:**
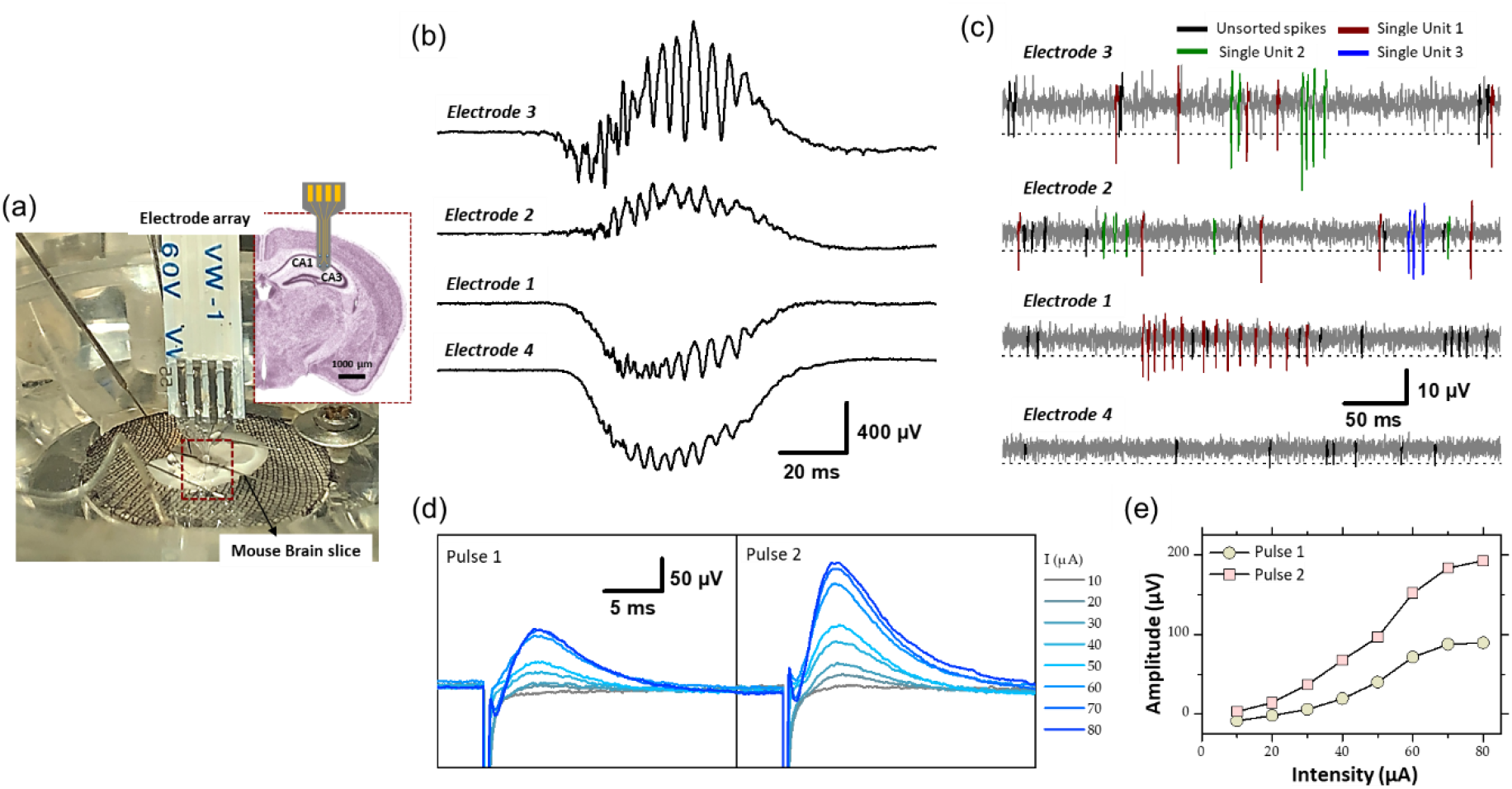
(a) Experimental set-up of the electrophysiological recording and stimulation, in the hippocampal region (CA1 and CA3) of a mouse brain slice, using flexible microelectrode array modified with PEDOT-CNF composite. Recording of spontaneous (b) sharp wave-ripples (filter: 0.1 Hz – 3 kHz) and (c) firing of action potentials from neurons (filter: 300 – 3000 Hz). Electrodes 3 and 2 were located in the pyramidal cell layer of CA1, and electrode 1 and 4 were in the dendrite layers. A threshold of −3xRMS (dashed line in figure 7c) was used for spike sorting. Single-units were identified using PCA and cluster analysis. Single-units are differentiated by color codes. Spikes in black correspond to unsorted (multiunit) activity. The traces in (b) and (c) were not obtained simultaneously but sequentially. (d) Evoked field potential recorded in the pyramidal cell layer after electrical stimulation of the Schaffer collateral. Pairs of cathodic square current pulses (200 μs width), with an interpulse interval of 20 ms, were applied at 0.5 Hz. Responses evoked by the first and second stimulus of the pairs are shown separately (pulse 1 and pulse 2). The different traces correspond to the response obtained at different stimulation intensities (10-80 μA). Each trace is the average of 10-20 responses obtained at a given intensity. (e) Field potential amplitude represented as a function of stimulation intensity. (For interpretation of the references to colours in this figure legend, the reader is referred to the Web version of this article).

## 3. Results and Discussion

### 3.1. Morphological study of electrodeposited PEDOT-CNF composites on flexible implants

In this work, we used a flexible neural implant having an array of 4 gold micro-disk-electrodes (40-μm diameter), that are sandwiched between two parylene C layers (Fig. 1a-b). A thick polymer backbone (23 μm) and thin passivation layers (1.3 μm) were opted for, such that there exists a balance between flexibility and improved long-term performance *vs* potential insulation regulation. We used this design to directly compare PEDOT-CNF and bare gold microelectrodes properties (structure, morphology, electrochemical performances and stability). The electrochemical deposition of PEDOT-CNF composite is illustrated in Fig. 1c-e. Before the electrodeposition, oxidized CNFs were synthesized for the following electrochemical synthesis of PEDOT-CNF composite [32, 61, 62]. Chemical oxidation of the CNFs leads to the formation of negatively-charged functional groups *i.e.* carboxylate and hydroxyl on the outer surface of the CNFs, which render them usable for charge-balancing anionic PEDOT dopant. Later, the PEDOT-CNF nanocomposites were deposited, on the flexible implantable electrode array, where PEDOT was galvanostatically deposited along with the entrapped oxidized CNFs within its matrix in one step (Fig. 1c-e). The deposition took place through a simultaneous oxidative PEDOT polymer chain propagation and CNF trapping mechanisms, resulting in a fibrous network of oxidized CNFs surrounded by PEDOT. The optimal deposition conditions were investigated by varying the surface charge densities, ranging from 1 nC/μm^2^ to 8 nC/μm^2^, at a constant current density of 10 pA/μm^2^.

SEM observations and FIB cross-sectional characterization of PEDOT-CNF composites of all deposition conditions (1 to 8 nC/μm^2^), showed that the entrapped CNFs were spatially distributed all over the gold electrode surface, and constitute a network of inter-connected nanofibers with variations in their aspect ratios and deposition densities (Fig. 2a-b). Since the CNFs core dictates the electrochemical growth of PEDOT, a porous and fibrous structure was a common feature among all the deposits. The deposition thickness and diameter, therefore the effective surface area, increased linearly with respect to the applied charge density (Fig. 2c-d). At the lowest charge densities (1 nC/μm^2^), the gold electrode surface was covered by a non-uniform layer of PEDOT-CNFs. In contrast, at higher charge deposition densities, the composite film was more uniformly porous and fibrous, thereby at the same time facilitating seamless intra- and interlayer ionic/electronic transport.

The energy dispersive X-ray (EDX) mapping data (Fig. 3) illustrates the spatial distribution of gold (white), sulfur (blue), carbon (red) and oxygen (green) within the gold-PEDOT-CNF composite electrode at a surface charge deposition density of 8 nC/μm^2^. Within the composite, the sulfur reflects the presence of PEDOT and the oxygen represents both PEDOT and COOH groups on oxidized CNFs. According to Fig. 3e, PEDOT is found to be densely concentrated at the gold interface, suggesting that there exists a highly ordered and less porous PEDOT-CNF composite at the gold-composite interface. In addition, the presence of sulfur was observed majorly around the walls of CNFs [32], indicating that PEDOT is grafted around the walls of nanofibers. Overall SEM and EDX observations suggest that PEDOT is acting as a polymer chain template to trap the oxidized CNFs and propagate all around the gold electrode surface resulting a three-dimensional growth of PEDOT around the oxidized CNFs.

### 3.2. Electrochemical characterization (CSCc, EIS and CIL)

The electrodeposited PEDOT-CNF composite films at different deposition charge densities were electrochemically characterized *in vitro* to assess their bidirectional transduction (electrolyte/electrode) capabilities. In this regard, electrochemical impedance spectroscopy (EIS), cathodic charge storage capacity (CSCc) and charge injection limits (CIL) are the essential parameters. On the one hand, a minimal impedance value is required to achieve signal noise reduction, such as thermal noise through shunt pathways [16, 18, 65]. On the other hand, large charge storage capacity and maximized charge injection limit values are particularly desired to establish safe electrical stimulation [18, 65].

#### 3.2.1. CSCc and EIS measurements

The evaluation of the charge transfer capabilities of PEDOT-CNF composites was carried out in aCSF, a physiologically relevant media, by sweeping a potential range between −0.85 V and 0.6 V, at a scan rate of 200 mV/s, using cyclic voltammetry (CV). This technique provides insights regarding the electrode interface under electrical load, the electrochemical conversion of species within the solution, and the transient changes due to redox reactions at the electrode surface [18]. The cathodal CSC (CSCc) of the composite film was calculated as the time integral of the cathodal currents within the cycled region. As indicated in Fig. 4a-b, the composite deposition on the gold microelectrode resulted in a progressive increment in the average CSCc values, up to 48 ± 4 mC/cm^2^, much higher than that of the bare gold electrode (1 ± 0.35 mC/cm^2^). There is a linear relationship between the charge delivered to the electrode during the deposition and charge storage capacity (Fig. 4b). This behavior is likely due to the surface area increment that allows for the effective diffusion of electrolyte ions at the electrode-solution interface.

The EIS measurements and phase angle responses of the composite electrodes were measured over a range of frequencies from 10 Hz to 7 MHz (Fig. 4c-f). The phase angle measurements of the PEDOT-CNF modified electrodes at lower frequency range (^~^10 Hz), revealed that the capacitive behavior was predominant with an angle around 80° (Fig. 4f). The angle shift towards resistive charge transfer, for a frequency range of 10 Hz to 10 kHz was proportional to the deposition charge density *i.e.* the higher the effective surface area, the larger the phase angle shift [57, 66]. Fig. 4c represents the bode plot displaying the impedance magnitude (|Z|) *vs.* frequency. PEDOT-CNF modified electrodes show an impedance range of 10-20 kΩ, which is at least 10 times less than that of the bare gold electrode (200-400 kΩ). Impedance values of the 5 deposition conditions were analyzed using one-way ANOVA followed by post-hoc Turkey’s test (n=5). ANOVA evidenced a significant effect of deposition charge density on |Z|_1 kHZ_ (*P* < 0.0001), yet changes in |Z|_1 kHZ_ were not proportional to the charge density, where the values at |Z|_1 kHZ_ being the most commonly used characteristic frequency band for action potentials [20]. In comparison to the 1 nC/μm^2^ charge density, significant lowering of |Z|_1 kHZ_ was only obtained with charge densities of 4 nC/μm^2^ and 6 nC/μm^2^ (*P* < 0.0001 and *P* = 0.0006 respectively, Tukey’s test). |Z|_1 kHZ_ obtained with charge densities of 2 and 8 nC/μm^2^ did not differ from that at 1 nC/μm^2^ (*P* = 0.3 and 0.06 respectively). The conditions 4 nC/μm^2^ and 6 nC/μm^2^ showed no statistically significant differences (P = 0.09). However, the optimal deposition condition seemed to be that at 6 nC/μm^2^ as it displayed both a sharp impedance decrement on average and the smallest variability across electrodes, with the |Z|_1 kHz_ value being 13.4 ± 2.2 kΩ (16.8 ± 2 MΩ.μm^2^).

To follow up further the investigation of the influence of the electrode surface area on the electrode performance, Nyquist plots were made to monitor the evolution of charge transfer resistance (semi-circle region) as a function of deposition charge on the electrode. As illustrated in Fig.4e the diameter of the semi-circle region becomes smaller with the increase in deposition charge, where it is governed by the electrode thickness, porosity and integrity of the Au-PEDOT-CNF composite interface [67, 68]. Among all the deposition conditions, 4 nC/μm^2^ and 6 nC/μm^2^ clearly showed depressed semi-circle regions, thus reflecting the improved area and highly conductive surface of the electrodes with lower impedance, possessing the capability of seamless bidirectional transfer of charges/electrons.

In the 8 nC/μm^2^ charge deposition case, even though the thickness and porosity of the electrode improved the effective surface area, it also probably induced mechanical stress at the Au-PEDOT-CNF interface, resulting in a slightly increased semi-circle region, therefore a charge transfer resistance and an impedance with larger variance compared to the 6 nC/μm^2^ condition. Overall, our results indicate that a charge density of 6 nC/μm^2^ optimized the deposition of PEDOT-CNF composite on the flexible gold electrode surface resulting in a specific impedance value of 16.8 ± 2 MΩ.μm^2^ at 1 kHz. This value is on par with the high-performance CNT fiber (20.5 MΩ.μm^2^) [69] and graphene fiber (9-28 MΩ.μm^2^) [70] microelectrodes, and 2 to 7 times lower, than most of the electrode materials deposited on flexible metallic substrates reported in the literature (Fig. 5f). The corresponding values and references are reported in Table S1.

#### 3.2.3. Electrical stimulation

As shown in the Fig. 4a, CSCc measurements with respect to the deposition charge density were used to identify the amount of charge available in the cathodic region of their respective CV sweep. Although CV at slow scan rate provides information related to the electrochemical reactions that occur at the electrode/electrolyte interface, it cannot reflect the amount of charge available during sub-millisecond stimulation pulses. Charge balanced square wave current pulses are generally used in electrical stimulation for electrophysiology experiments and therapies, with pulse widths ranging from 50 to 1000 μs. A high CIL would be beneficial for these extremely fast charge-discharge processes by preventing damages to the tissue-electrode region by reducing the incidence of irreversible Faradic reactions [18, 20]. To assess the charge stimulation capability of the PEDOT-CNF composite, as a prerequisite, identifying the water electrolysis limits, especially the water reduction limit in physiologically relevant solution is important. Fig. 5a shows that the water reduction voltage of PEDOT-CNF modified gold electrodes is around −1.3 V. Next, the voltage excursions in response to biphasic, cathodic first, current pulses were recorded with a 1000 μs pulse width in aCSF (Fig. 5b-d). Using a range of pulse current intensities, we defined the CIL as the amount of charge injected which caused polarization (V_p_) of the electrode beyond its water hydrolysis window. From the lowest to highest deposition charge density, the CIL values increased linearly up to the condition 6 nC/μm^2^, being the best one among all, with an average value of 7.6 ± 1.3 mC/cm^2^ (Fig. 5e). This result agreed with the corresponding impedance measurements where the 6 nC/μm^2^ deposition condition provided a low impedance value with lowest variability (Fig. 4d).

The optimized CIL value from this work was compared with other PEDOT based composites [40, 71, 72] and porous metals deposits [69, 73–76] on flexible neural implants. Fig. 5f and Table S1 shows that, among all reported electrode materials deposited on flexible metallic substrates, the PEDOT-CNF coating displayed superior electrical properties and charge injection capabilities, thanks to the chemical composition of the material and its physical morphology.

#### 3.2.3. Long time performance

Along with the charge injection capability, it was also important to assess the biphasic charge cycling endurance. For this purpose, the electrodes were subjected to a series of charge balanced current pulses in physiologically relevant aCSF. Our electrodes were repeatedly pulsed over 0.5 million times using the 200 μs cathodic pulse width, followed by an immediate charge-compensating anodic pulse (most commonly reported) [20], consisting in a charge injection density of 0.5 mC/cm^2^ (current amplitude of 33 μA). Fig. 6a shows the evolution of the polarization potential (V_p_) during the time of charge injection pulsing. A resultant V_p_ drift of 0.3 V, from −0.1 V to −0.4 V, was observed after 0.5 million pulses, but remained far from the water window critical limit (−1.3 V) throughout the stimulation. The corresponding |Z|_1 kHz_ value was increased from an initial value of 18 kΩ to a-final value of 38 kΩ after 0.5 million biphasic stimulation cycles, while still being far from the bare non-coated Au electrode |Z|_1 kHz_ value (>200 kΩ) (Fig. 6b). In addition, the SEM images of the composite after the stimulation *vs* control, from the Fig. 6c, suggested that the electrodes exhibited no significant physical delamination or degradation even after 0.5 million biphasic stimulation cycles confirming an excellent structural control and longevity.

### 3.3. Brain slice electrophysiological recording and stimulation

Electrophysiological experiments were conducted to assess the recording quality of PEDOT-CNF electrodes and their usability as stimulating electrodes (Fig. 7a). Electrical stimulation and recordings were performed in the hippocampus (CA1 and CA3 regions) of mouse brain slice maintained *in vitro* [77]. PEDOT-CNF modified electrodes allowed to record two types of spontaneous neuronal activities: sharp wave-ripples (SWR) complexes and neuronal spiking activity.

SWR complexes constitute a mesoscopic signal that reflects synchronized activity in large population of neurons [77]. SWR complexes were recorded with minimal filtering (0.1 Hz-3 KHz) where two electrodes of the flexible four electrode probe were located in the cell layer of CA1 and other two in the dendrite layer (stratum radiatum/lacumosum). As shown in the Fig. 7b, the slow component of the SWR complex, “the sharp wave” proper, shows an inversion in polarity between the two regions: positive in the cell layer (electrodes 3 and 2) and negative in the dendrite layer (electrodes 1 and 4), as expected given the fact that the excitatory synaptic inputs, which generate the sharp wave are located on the dendrite of hippocampal pyramidal cells. The ripples correspond to the oscillatory pattern (200-250 Hz) carried by the sharp wave. The amplitude of ripples was larger in the cell body layer (electrode 3 and 2), as expected since they mostly correspond to the population spikes patterned by the local inhibitory neurons.

The other type of spontaneous activity that was recorded in the hippocampus corresponds to action potentials APs (spikes) generated by neurons [77, 78]. They were visualized between sharp wave-ripple complexes while using bandpass filtering (300 - 3000 Hz). The examples presented in Fig. 7c, were obtained with the same PEDOT-CNF modified electrodes and at the same electrode locations as in Fig. 7b. The APs are recognized as fast and mostly negative deflections of varying amplitude. To proceed further we examined whether single-unit activity, i.e., APs that can be attributed to one single neuron, could be extracted from the raw traces. First portions of trance around events were extracted using a threshold at 3xRMS of the voltage trace (dashed lines in Fig. 7c), and thereafter analyzed by PCA and clustering (not illustrated). This analysis allowed identifying constant spike shapes, which were further ascribed to single-unit activities if the inter-spike interval distribution (not illustrated) showed a clear refractory period, i.e., no interval <1 ms. In contrast, multiunit activities (black spikes in Fig. 7c) do not have a refractory period because they are produced by several independent neurons. As shown in Fig. 7c, two and three single-units (color coded) have been isolated from electrodes 3 and 2, respectively and only one single-unit was isolated with electrode 1. This is likely due to the fact that only few neuronal cell bodies, from which action potentials are generated, are to be found in the dendrite layers of the hippocampus. In summary, the resulting signal-to-noise ratio of the PEDOT-CNF electrodes facilitated the isolation of singleunit activity from neighboring neurons.

In addition, the PEDOT-CNF electrodes were validated for their usability as stimulating electrodes. Electrical stimulation was applied through a PEDOT-CNF electrode positioned in the dendritic layer (stratum radiatum) to activate axons of the Schaffer collateral (the projection issued from CA3 pyramidal cells and synapsing on the dendrites of CA1 pyramidal cells) (Fig. 7d-e). The extracellular recording of the local field potential was made in the cell body layer of CA1 with a conventional metallic electrode. Electrical stimulus were given in pairs (20 ms interpulse interval) and, consisted in cathodic square current pulses (200 μs) delivered at different intensities. Pulse pairs were repeated at a frequency of 0.5 Hz to examine short-term plasticity of the synaptic responses. For suprathreshold intensities, the potential was positive due to recording in the cell layer (current sink in the dendrites), and the amplitude of the response to the second pulse of the pair was larger than that of the first one, reflecting short-term facilitation of the synaptic response (Fig. 7d). The amplitude of the potential increased in proportion to the stimulation intensity, with a saturation at around 80 μA (Fig. 7c). This saturation suggests that all the axons implicated in the response were recruited with an intensity ≤ 80 μA. Thus, PEDOT-CNF microelectrodes could be used as efficient stimulating electrodes, to obtain typical evoked potentials with charge densities that remained below the water reduction limit.

## 4. Conclusion

We have developed a well-controlled and versatile surface modification method for preparing macroporous PEDOT-CNFs microelectrodes on flexible implantable neural probes. We investigated by FIB and EDX the mechanism by which a macroporous nanostructure of PEDOT-CNF layer was created, where the conducting PEDOT polymer was covered uniformly and tightly around the oxidized carbon nanofibers as a solid doping template. We found that the combination of carbon nanofibers and PEDOT as a single carbonaceous composite resulted in a strong synergetic effect leading to a lower impedance, superior charge storage capacity and charge injection limit compared to bare metal and other reported organic coated flexible electrodes in electrochemical characterization. We further showed that that the carbon nanofibers function as reinforcing elements within the composite material and prevent PEDOT film from undergoing delamination or cracking during long lasting electrical pulsing experiments. *In vitro* experiments on mouse brain slices showed that the PEDOT-CNFs microelectrodes can record spontaneous oscillatory field potentials as well as single-unit action potentials with good signal-to-noise ratio, and allow to safely deliver electrical stimulation for evoking field potentials. These results show that these electrodes are well suited for high-performance recording and/or stimulation for applications in brain therapies.

## CRediT authorship contribution statement

**Venkata Suresh Vajrala:** Methodology, Investigation, Formal analysis, Visualization, Validation, Writing-original draft & editing. **Valentin Saunier:** Investigation, Methodology, Writing - review & editing. **Lionel Nowak:** Conceptualization, Investigation, Formal analysis, Writing-review & editing. **Emmanuel Flahaut:** Conceptualization, Writing - review & editing. **Christian Bergaud:** Supervision, Resources, Conceptualization, Writing-review & editing. **Ali Maziz:** Conceptualization, Supervision, Project administration, Resources, Methodology, Formal analysis, Writing-original draft & editing.

## Declaration of competing interest

The authors declare that they have no known competing financial interests or personal relationships that could have appeared to influence the work reported in this paper.

## Acknowledgment

This project was financially supported by the CNRS (“Centre National de la Recherche Scientifique”) and the ANR (“Agence Nationale pour la Recherche”, project 3D-Brain (ANR-19-CE19-0002-01). The technological realisations and associated research works were partly supported by the French RENATECH network.

## Appendix A. Supplementary data

The following is the Supplementary data to this article:

---

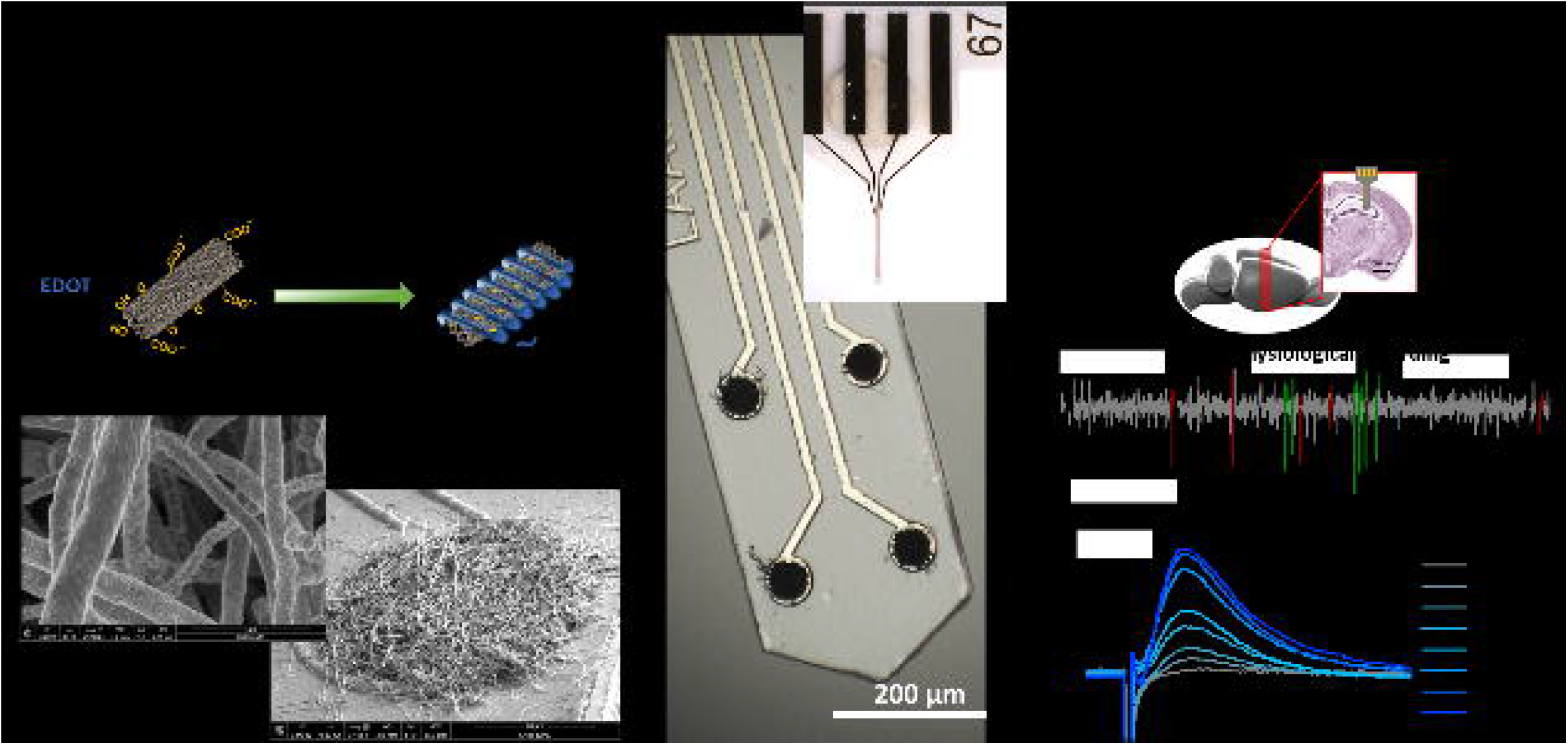

## Notes

### Competing Interest Statement

The authors have declared no competing interest.

